# Concentration of 2’C-methyladenosine triphosphate by *Leishmania guyanensis* enables specific inhibition of *Leishmania* RNA virus 1 via its RNA polymerase

**DOI:** 10.1101/261172

**Authors:** John I. Robinson, Stephen M. Beverley

## Abstract

*Leishmania* is a widespread trypanosomatid protozoan parasite causing significant morbidity and mortality in humans. The endobiont dsRNA virus *Leishmania* RNA virus 1 (LRV1) chronically infects some strains, where it increases parasite numbers and virulence in murine leishmaniasis models, and correlates with increased treatment failure in human disease. Previously, we reported that 2’-C-methyladenosine (2CMA) potently inhibited LRV1 in *Leishmania guyanensis* (*Lgy*) and *L. braziliensis*, leading to viral eradication at concentrations above 10 µM. Here we probed the cellular mechanisms of 2CMA inhibition, involving metabolism, accumulation and inhibition of the viral RNA dependent RNA polymerase (RDRP). Activation to 2CMA triphosphate (2CMATP) was required, as 2CMA showed no inhibition of RDRP activity from virions purified on cesium chloride gradients. In contrast, 2CMA-TP showed IC50s ranging from 150 to 910 µM, depending on the CsCl density of the virion (empty, ssRNA- and dsRNA-containing). *Lgy* parasites incubated *in vitro* with 10 µM 2CMA accumulated 2CMA-TP to 410 µM, greater than the most sensitive RDRP IC50 measured. Quantitative modeling showed good agreement between the degree of LRV1 RDRP inhibition and LRV1 levels. These results establish that 2CMA activity is due to its conversion to 2CMA-TP, which accumulates to levels that inhibit RDRP and cause LRV1 loss. This attests to the impact of the Leishmania purine uptake and metabolism pathways, which allow even a weak RDRP inhibitor to effectively eradicate LRV1 at micromolar concentrations. Future RDRP inhibitors with increased potency may have potential therapeutic applications for ameliorating the increased Leishmania pathogenicity conferred by LRV1.

## Introduction

The neglected tropical disease leishmaniasis is caused by species of the genus *Leishmania*, which are single-celled eukaryotic parasites transmitted by phlebotomine sand flies (1,2). Leishmaniasis occurs in many regions of the world, with more than 12 million cases and more than 1.7 billion people at risk (1,3). Three clinical presentations are most common: mild, self-healing cutaneous lesions (CL), fatal visceral disease, and disfiguring metastatic forms such as mucocutaneous leishmaniasis (MCL) (4). The factors determining disease progression and responsiveness to treatment are unclear, but are thought to be both host- and pathogen-derived. (5,6).

Many isolates of *Leishmania* within the subgenus *Viannia*, including *L. braziliensis (Lbr)* and *L. guyanensis (Lgy)*, bear *Leishmaniavirus*, a single-segmented dsRNA totivirus known as *Leishmania* RNA virus 1 (LRV1) ^1^ (5,7-9). Like most other Totivirus species, LRV1 is neither shed nor infectious and is inherited vertically (10,11); indeed, phylogenetic evidence suggests that LRV1 strains have persisted and co-evolved with their *Leishmania* hosts over millions of years (10).

Previous work has established that mice infected with parasites containing the endobiont LRV1 exhibit greater pathology, higher parasite numbers, and increased metastasis (12,13). These studies benefited from the availability of isogenic LRV1+ or LRV1- lines, generated spontaneously or by defined methods such as RNA interference or antiviral drug treatment (14-16).

The role of LRV1 in human leishmaniasis has been more challenging to establish definitively. When comparing rates of CL and MCL, some studies find that LRV1+ strains generate more MCL (17-19), while others do not (20,21). These discrepant findings may be explained by other parasite or host factors known to contribute to MCL pathology (13,22,23). Furthermore, differences in the severity of disease are not always accurately captured by binary categorization as CL or MCL. Moreover, co-infections with viruses inducing Type I interferon responses exacerbate *Lgy* pathology and metastasis (24,25), potentially obscuring the contributions of LRV1. Importantly, the presence of LRV1 in clinical isolates of *Lbr* or *Lgy* correlates with drug-treatment failure and relapses (18,20), which could be explained by the increased parasite numbers or altered host responses predicted from animal models (12,13,26). Overall, there is good reason to postulate a role for LRV1 in increasing disease severity in human leishmaniasis (13), although many questions remain.

LRV1 follows a typical totivirus life cycle where the dsRNA viral genome encodes two large overlapping reading frames, the capsid and RNA-dependent RNA polymerase (RDRP) (Fig. 1A). First, the dsRNA genome is transcribed by the viral RDRP into positive-sense ssRNA. As in many totiviruses, the RDRP is translated via a +1 frameshift, generating a capsid-RDRP fusion (27-29). The capsid monomers then self-assemble into immature virions (30), incorporating the positive-sense ssRNA transcript, which the RDRP then replicates into the mature dsRNA genome.

**Figure 1.**
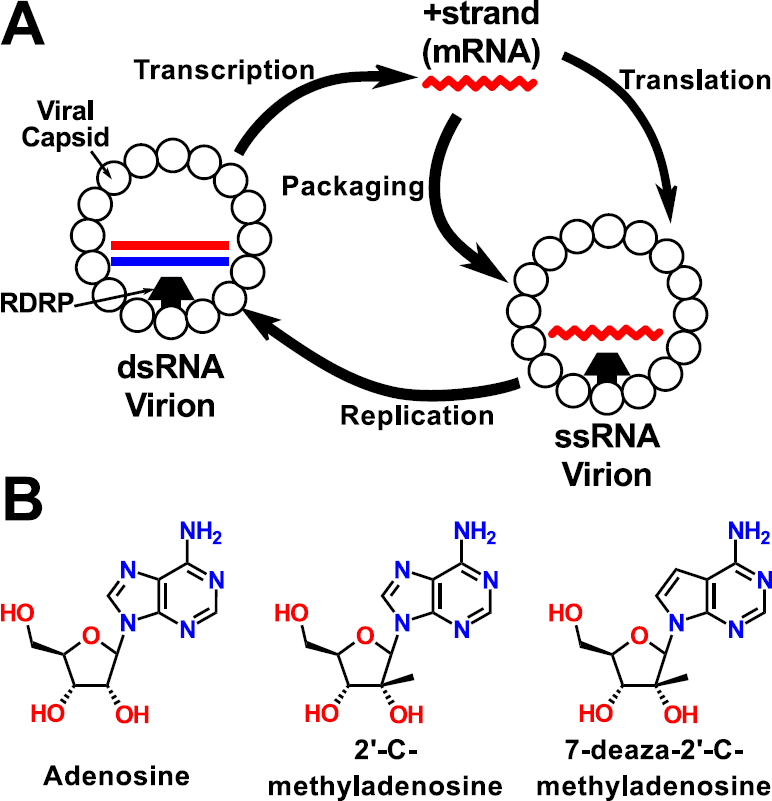
*Leishmania* LRV1 replication cycle and inhibitors. **Panel A.** A schematic depiction of the LRV1 lifecycle. RNAs are indicated in color (+ strand red, -strand blue); the dsRNA genome within the mature virion is shown as straight lines, while ssRNAs are shown as jagged lines. The viral RDRP (black trapezoid) is shown fused to a capsid monomer (white circle), as the RDRP is generated through frame shift translation (27-29). **Panel B.** Chemical structures of adenosine, 2’-C-methyladenosine, and 7-deaza-2’-C-methyladenosine.

Vaccination of mice using the LRV1 capsid results in significant protection against LRV1+ *Lgy* (31), suggesting that therapies targeting LRV1 specifically might aid in reducing disease pathology. Previously, we reasoned that the powerful nucleoside and nucleobase salvage pathways of *Leishmania* might enhance the efficacy of nucleosides analogs targeting the viral RDRP (15,32). Accordingly, screening a small library of antiviral nucleosides identified two closely-related adenosine analogs, 2’-C-methyl adenosine (2CMA) and 7-deaza-2’-C- methyl adenosine (7d2CMA) (Fig. 1B), which specifically inhibited LRV1 replication in cultured *Leishmania* cells (15). These compounds exhibited EC50s of 3-5 µM for viral inhibition, contrasting with much greater EC50s for the parasites themselves. The active compounds rapidly eradicated LRV1 when tested at concentrations above 10 µM, allowing us to readily create isogenic LRV1- lines (15).

Importantly, these were the first studies showing specific inhibition of any totivirus. The mechanism of anti-LRV1 activity was postulated to be through direct inhibition of the LRV1 RDRP by the triphosphorylated form of 2CMA. Here we provide support for this hypothesis, although the potency of 2CMA-TP for viral inhibition was unexpectedly weak. Remarkably, viral inhibition was accomplished through hyper-accumulation and retention of 2CMA-TP, arising from the powerful uptake and metabolic salvage pathways of these purine auxotrophs (32). These findings have significant implications for future efforts aimed towards developing new and more potent *Leishmania* virus inhibitors.

## Results

### Purification and separation of virion populations on CsCl gradients

RDRP assays were carried out with *Lgy* strain M4147 LRV1 virions purified by CsCl equilibrium gradient centrifugation (7,33). After fractionation, virions were detected and quantified by their reactivity with an anti-capsid antibody. We reproducibly observed three overlapping ‘peaks’, designated low-, medium-, and high-density (LD, MD and HD) (Fig. 2). In previous studies of the yeast L-A Totivirus, similar peaks were shown to correspond to virions that were primarily ‘empty’ or contained ssRNA or dsRNA, respectively (34,35). The densities of the *Lgy* LRV1 LD, MD and HD peaks were 1.29, 1.36 and 1.41 g/mL, in good agreement with the densities of L-A virus particles bearing ssRNA- and dsRNA- (1.31 and 1.41 g/mL, respectively) (36). Preliminary data from S1 nuclease digestion of viral RNA from these fractions were consistent with these assignments^2^.

**Figure 2.**
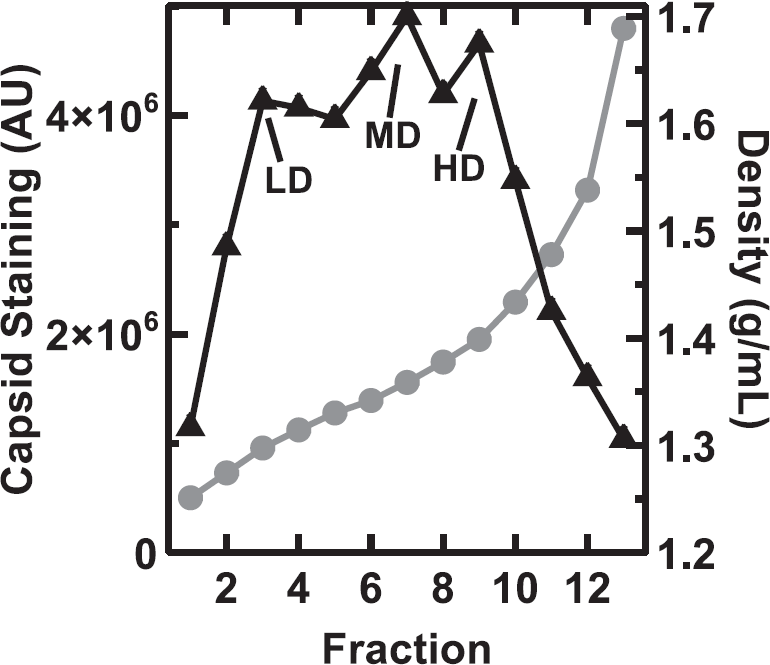
Distribution of viral capsid protein across a CsCl density gradient. Clarified parasite lysates were separated on a CsCl density gradient and the relative amounts of viral capsid protein in each fraction were measured (black triangles; see “Experimental Procedures”). The density of each fraction (gray circles) was measured with an Abbe refractometer. Data for one representative gradient are shown out of the 7 performed. The low, medium and high density “peak” fractions (LD, MD and HD) taken for RDRP assays, are labeled.

### *In vitro* assay of LRV1 RDRP activity

To measure RDRP activity, purified virions were allowed to incorporate [α-^32^P]UTP in the presence of the other three nucleoside triphosphates for 1 hr, a time sufficient for one round of viral genome replication (33). RNA was purified and separated by native gel electrophoresis, and the products were visualized and quantified. Two products were always found: one about 5 kb, presumably corresponding to the full-length LRV1 genome, and smaller, heterogeneous products ranging from 0.1 – 0.5 kb, which we considered abortive transcripts (Fig. 3A). Neither extending the incubation time nor increasing the concentration of UTP significantly altered the profile obtained^2^. Importantly, neither full-length nor small products were produced by corresponding preparations from LRV1-negative parasites (Fig. 3B).

**Figure 3.**
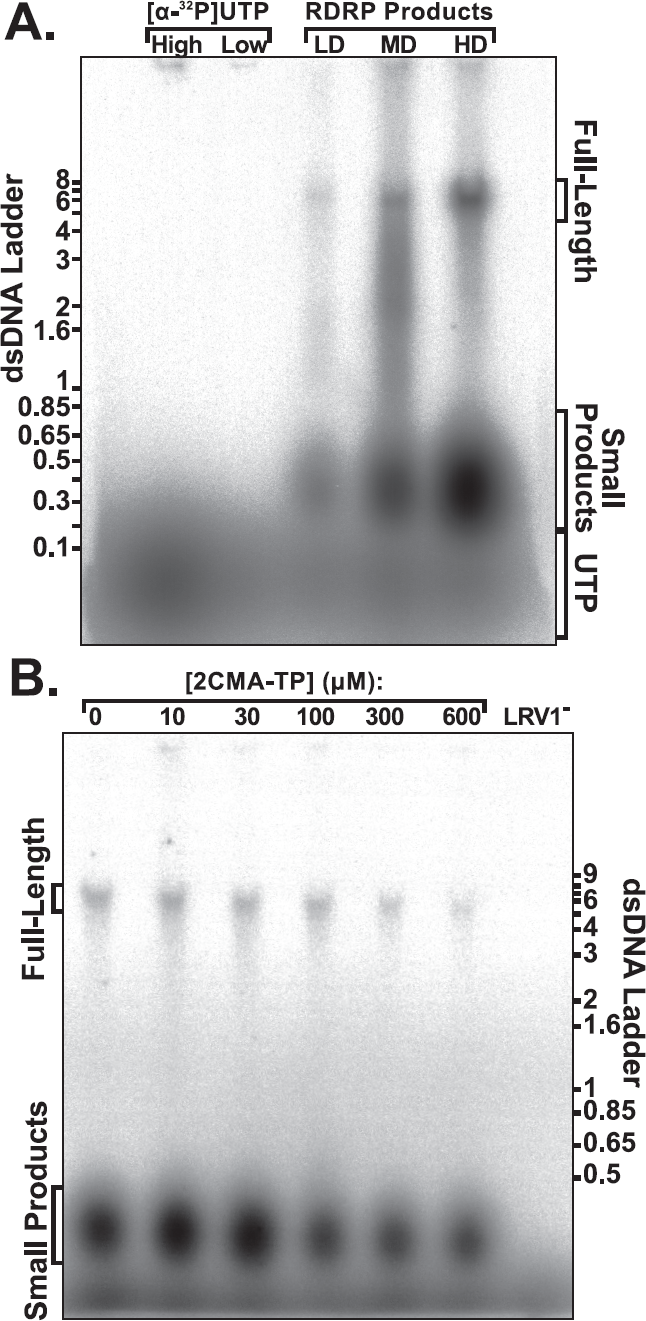
Radiolabeled RNAs produced by purified *Lgy* LRV1 RDRP *in vitro*. **Panel A.** RDRP activity in LD, MD, and HD fractions (Fig. 2) was assayed by [α-^32^P]UTP incorporation,as described in “Experimental Procedures.” Radiolabeled RNAs were run along-side pure [α-^32^P]UTP on a native agarose gel. The full-length and small RDRP products are labeled for reference. **Panel B.** RDRP reactions were performed in the presence of 0, 10, 30, 100, 300, or 600 µM 2CMA-TP. As a negative control, the RDRP reaction was run using a mock HD fraction isolated from LRV1- *Lgy* parasites. A representative titration using the HD fraction is shown here out of the 4 HD fraction titrations performed.

### 2CMA-TP specifically inhibits viral RDRP activity

Incubation of the three LRV1 fractions with 2CMA-TP reduced synthesis of both the full-length and small RDRP products (Figs. 3 - 5). The synthesis of each product was quantitated and normalized to that obtained with drug-free controls. IC50s were calculated by fitting the inhibition data with a logistic dose-response curve (Table 1). These ranged from 150 μM for full-length product synthesis by LD virions to 910 µM for small products synthesized by HD virions (Table 1, Figs. 3, 4). These IC50s were unexpectedly high, greatly exceeding the extracellular concentration of 2CMA shown previously to cause 50% inhibition of LRV1 abundance (˜3 µM) (15). This did not arise artificially from 2CMA-TP degradation during the assay, as HPLC tests of RDRP reactions showed only 3.7 ± 0.6% (n = 3) loss of 2CMA-TP.

**Figure 4.**
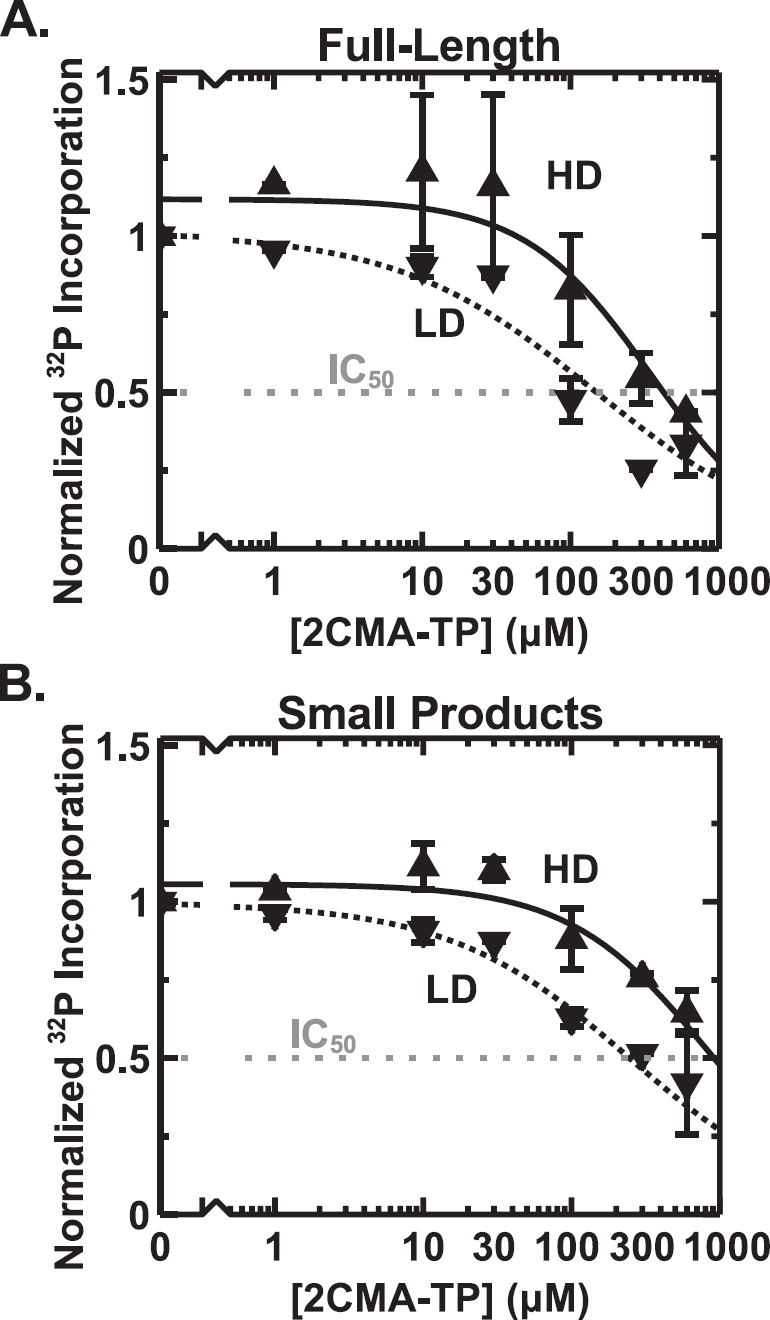
2CMA-TP Inhibition of RDRP activity of purified *Lgy* LRV1 virions. The amounts of full-length or small RDRP products were quantified and then normalized to the amount of product formed in the absence of 2CMA-TP. The averages and SDs (calculated with Microsoft Excel) from three LD virion titrations (dotted line) and four HD virion titrations (solid line) are shown. **Panel A.** Plot of full-length RDRP products. **Panel B**. Plot of small RDRP products. MD virions showed intermediate profiles^2^ (Table 1).

**Table 1.** Effect of 2CMA-TP on *Lgy* LRV1 RDRP activity.

| **Product** | **LRV1 fraction IC50 (μM)** |  |  |
| --- | --- | --- | --- |
|  | **Low Density**<br>**(n=3)** | **Medium Density**<br>**(n=3)** | **High Density**<br>**(n=4)** |
| Full-Length | 150 (90 – 210) | 250 (100 – 400) | 420 (310 – 540) |
| Small | 250 (160 – 330) | 640 (300 – 980) | 910 (620 – 1200) |

As anticipated, 2CMA did not measurably inhibit RDRP activity when tested at concentrations up to 1000 µM (Fig. 5). Similarly, dATP, which lacks both the 2’-hydroxyl and methyl groups of 2CMA (Fig. 1B), failed to inhibit RDRP activity at the highest concentration tested (600 µM; Fig. 5); nor in limited tests did 600-1000 µM TTP inhibit^2^. These data show that high concentration of a non-substrate nucleoside triphosphate do not inhibit RDRP activity nonspecifically. Together, these data suggest that despite its relatively low potency, under these assay conditions 2CMA-TP inhibition of LRV1 RDRP activity was specific.

**Figure 5.**
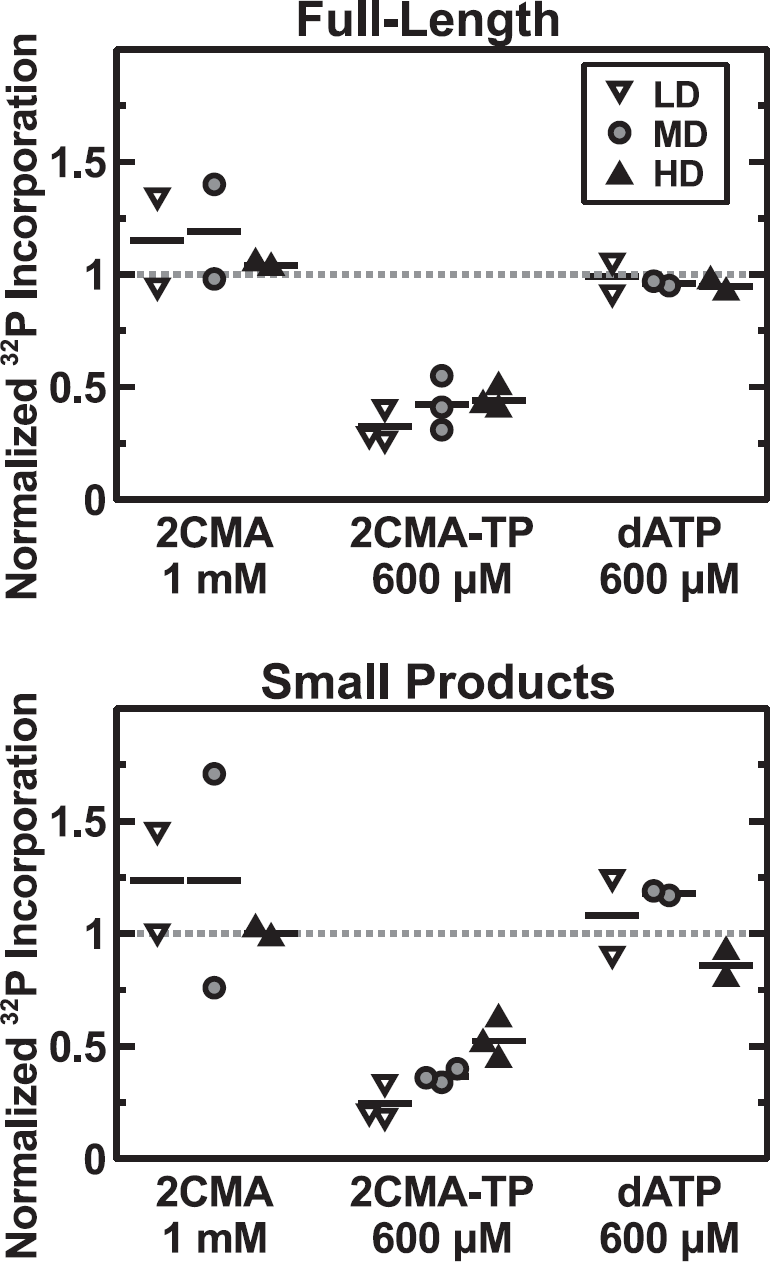
Specificity of *Lgy* RDRP inhibition by 2CMA-TP relative to 2CMA and dATP. RDRP reactions were performed for 1 hour in the presence of 1 mM 2CMA, 600 µM dATP, or 600 µM 2CMA-TP. The amount of full-length (top panel) and small products (bottom panel) were measured and normalized to untreated control reactions. Individual values are shown for two experiments (with 1 or 2 technical replicas) and the means are marked with horizontal lines.

### 2CMA activation to 2CMA-TP within parasites

To account for the relative sensitivity of *Lgy* LRV1 within cells to 2CMA treatment compared to the insensitivity of LRV1 RDRP activity to inhibition by 2CMA-TP, we hypothesized that parasites must acquire and activate 2CMA, accumulating high 2CMA-TP concentrations. We developed an HPLC protocol capable of resolving synthetic 2CMA-TP from natural ribonucleotides and the internal standard, dGTP (Fig. 6A-B; a small peak presumed to be 2CMA-DP was also observed). Although dGTP occurs naturally, its intracellular concentration of approximately 5 µM is well below the limit of detection for this assay and thus does not interfere with its use for this purpose.(37). Standard mixtures were then used to generate a calibration curve relating peak area to 2CMA-TP amount.

**Figure 6.**
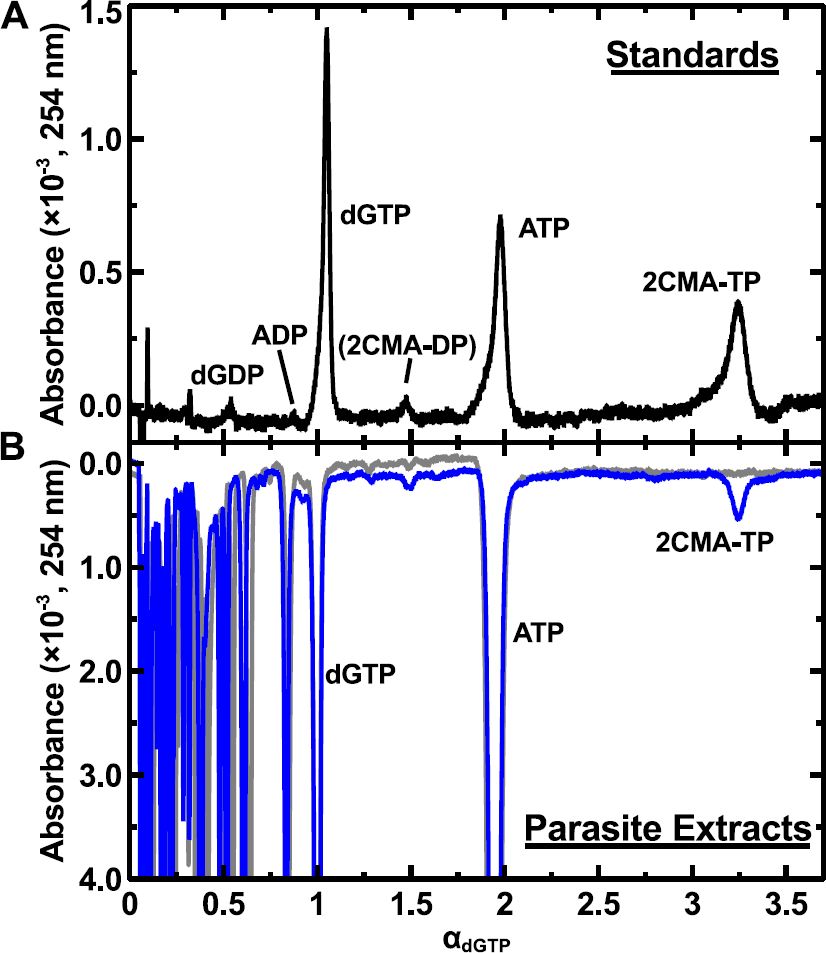
*Lgy* M4147 LRV1+ parasites synthesize 2CMA-TP. **Panel A.** Standard mixture of dGTP, ATP, and 2CMA-TP establishing the HPLC elution profile of 2CMA-TP relative to dGTP, the exogenous internal standard (endogenous cellular dGTP concentrations are well below the limit of detection). The small peak eluting between dGTP and ATP is presumed to be 2CMA-DP. **Panel B.** Detection of 2CMA-TP in *Lgy* incubated with 10 µM 2CMA for 20 hours (blue line). An extract from cells grown without drug (gray line) is provided for comparison. In order to correct for variation in extraction efficiency and HPLC elution times, *Lgy* samples had 7 nmol dGTP spiked in immediately prior to extraction.

We next evaluated several protocols for extracting parasite nucleotides and determined that extraction with 1:1 acetonitrile:water performed best, as judged by recovery of the added dGTP standard (Experimental Procedures). We then compared the nucleotide profiles of LRV1+ *Lgy* grown in the presence or absence of 10 µM extracellular 2CMA for 20 hours, a time corresponding to more than two rounds of parasite replication. Under these conditions, we observed a peak co-eluting with synthetic 2CMA-TP that was absent from untreated parasites (Fig. 6B). This established the parasite’s capacity to activate 2CMA to 2CMA-TP.

### Time and external 2CMA concentration dependence of 2CMA-TP accumulation

*Leishmania* cell volumes vary somewhat depending on the culture’s growth phase (38). To calculate the intracellular 2CMA-TP concentration, we therefore determined the average volume of *Lgy* parasites under our assay conditions. In logarithmic growth phase, parasite volumes averaged about 23 fL (Experimental Procedures).

We measured the intracellular concentration of 2CMA-TP as a function of time of incubation with 10 μM external 2CMA. Within 1.5 hours, 2CMA-TP levels had risen to concentrations comparable to the minimal RDRP IC50 levels (Fig. 7A), and had reached maximal levels by 8 hours.

**Figure 7.**
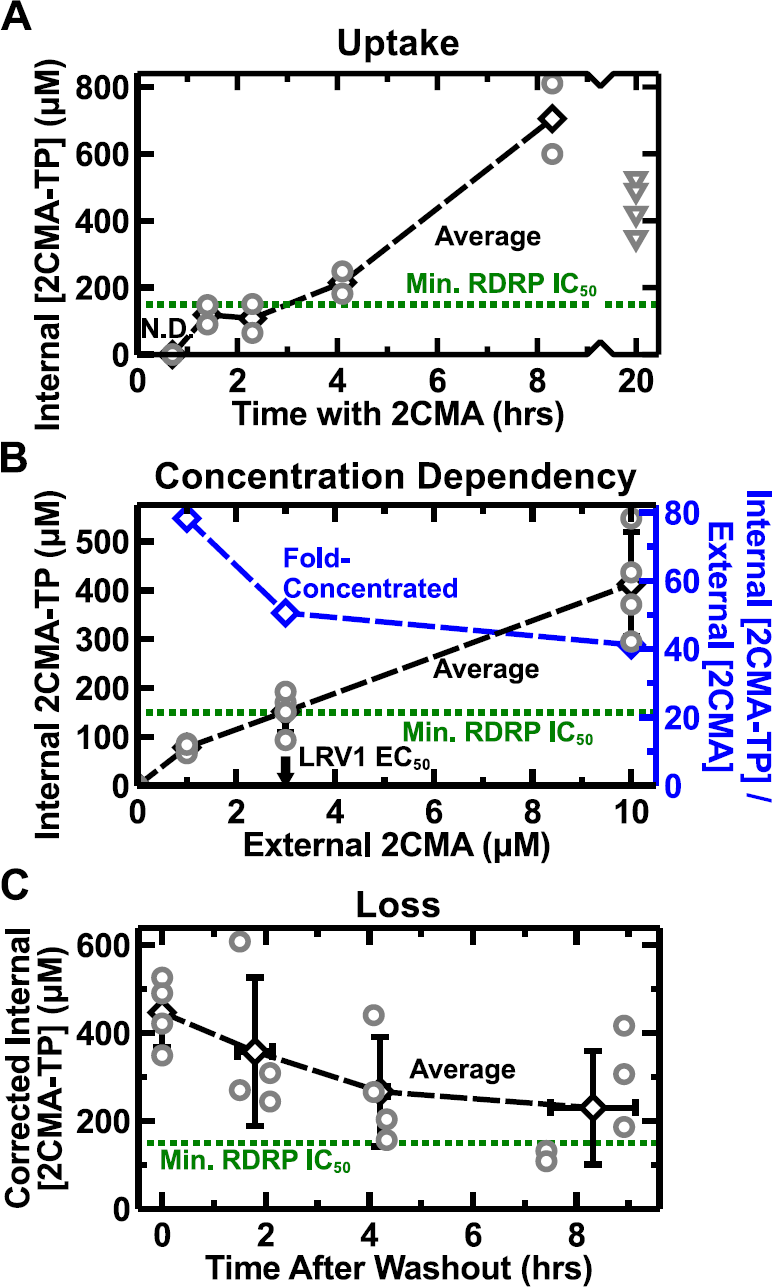
*L. guyanensis* parasites accumulate high levels of 2CMA-TP. Intracellular 2CMATP was extracted and quantified by HPLC (Fig. 6; Fig. S2) and its concentration was calculated using the measured cell volume of 23 fL under these conditions (Experimental Procedures). Each replicate (grey circles) was obtained from a separate 5-mL culture grown in Schneider’s medium. Averages and standard deviations (black diamonds; calculated in Microsoft Excel) are also plotted for each time point or condition. For reference, the minimum RDRP IC50 is marked on each graph (green dashed line). **Panel A.** 2CMA-TP accumulation in parasites measured over time. *Lgy* parasites were incubated in 10 µM 2CMA for the indicated periods and 10^8^ cells were harvested for analysis. This experiment was performed twice. Intracellular 2CMA-TP concentrations at equilibrium after 20 hours of incubation with 10 µM 2CMA are also plotted(grey triangles). **Panel B**. Steady-state concentrations of 2CMA-TP vary with 2CMA dose. *Lgy* parasites were grown for 20 hours in the presence of the indicated concentrations of external 2CMA prior to analysis. This experiment was repeated four times. The fold-concentration of intracellular 2CMA-TP was calculated relative to extracellular 2CMA dose (blue diamonds). For reference, the EC50 for LRV1 inhibition in parasites is marked on the X axis (black arrow). **Panel C**. Retention of 2CMA-TP following removal of 2CMA. LRV1+ *Lgy* M4147 parasites were incubated for 20 hours in the presence of 10 µM 2CMA; at that time, cells were harvested, washed, and resuspended in drug-free medium. Intracellular 2CMA-TP levels were determined after incubation for the indicated times. To correct for 2CMA-TP dilution due to parasite replication, 2CMA-TP concentrations were multiplied by the fold population growth since 2CMA removal (about 1.4-fold over the course of the experiment). This experiment was repeated twice using two independent cultures per time point (n=4; grey circles). Averages and standard deviations (black diamonds; calculated in Microsoft Excel) are also plotted for each time point. Since some time points were collected at slightly varying times, these were averaged for plotting in the figure.

We then measured the intracellular 2CMA-TP levels following growth in varying levels of 2CMA for 20 hours (Fig. 7B). The concentrations were chosen to span a range from non-inhibitory (1 μM) to that sufficient to eradicate LRV1 entirely (10 µM) (15). Over this range of concentrations, internal 2CMA-TP levels exceeded external 2CMA concentrations by 40- to 80-fold (Fig. 7B).

When propagated in the sub-inhibitory dose of 2CMA (1 µM), the intracellular 2CMA-TP concentration was 78 ± 9 μM (Fig. 7B, n=4), well below the minimal IC50 for RDRP inhibition (150 µM for full-length products from LD virions; Table 1). When treated with 3 µM external 2CMA—the EC50 for LRV1 inhibition— intracellular 2CMA-TP levels rose to 152 ± 43 μM (n = 4), comparable to the minimal RDRP IC50. Finally, cells propagated in 10 µM external 2CMA, reached an intracellular 2CMA-TP concentration of 410 ± 110 μM (n=4) (Fig. 7B), well in excess of the minimum IC50 for RDRP inhibition.

Thus, with increasing external 2CMA, the steady-state levels of intracellular 2CMA-TP rose progressively to values exceeding the minimal RDRP IC50, which in turn corresponded reasonably well to the observed effects on LRV1 inhibition.

### Parasites retain 2CMA-TP for an extended period following 2CMA removal

Once phosphorylated by cells, some nucleoside analogs are retained for an extended period as triphosphates, thereby contributing significantly to drug efficacy (39). To assess this, *Lgy* parasites were incubated for 20 hours in 10 µM 2CMA, washed, and resuspended in drug-free medium, after which intracellular 2CMA-TP levels were measured and corrected for parasite replication (Fig. 7C). Even after 8 hours, 2CMA-TP levels remained at or above the minimal RDRP IC50. Indeed, the persistence of intracellular parasite 2CMA-TP greatly exceeded that of serum 2CMA measured in mammalian models (40).

### A quantitative simulation of RDRP and virus inhibition

We asked whether a simple computational model using the relative rates of parasite and viral replication under conditions of drug treatment could quantitatively simulate the experimental data above. We developed a model employing Gibson and Bruck’s next-reaction modification to Gillespie’s stochastic simulation algorithm (41), which directly simulates the occurrence of individual events over time by picking the next event-time from an associated probability distribution (Experimental Procedures; Supplemental File 2).

In this model, the relevant parameters were parasite and virus replication rates. We used the experimentally-measured parasite population doubling time of 7.5 hours and assumed that in the absence of drug pressure the relative parasite and virus replication rates were identical. Simulations began with an initial population of 1000 cells, a number large enough that repeated simulations deviated <1% from each other. The simulated parasites each initially contained 16 LRV1 particles, consistent with prior measurements of virus numbers (15). Using these conditions, the simulation correctly maintained the LRV1 population over time at an average of 16 virions per parasite in the absence of drug pressure (Fig. 8).

**Figure 8.**
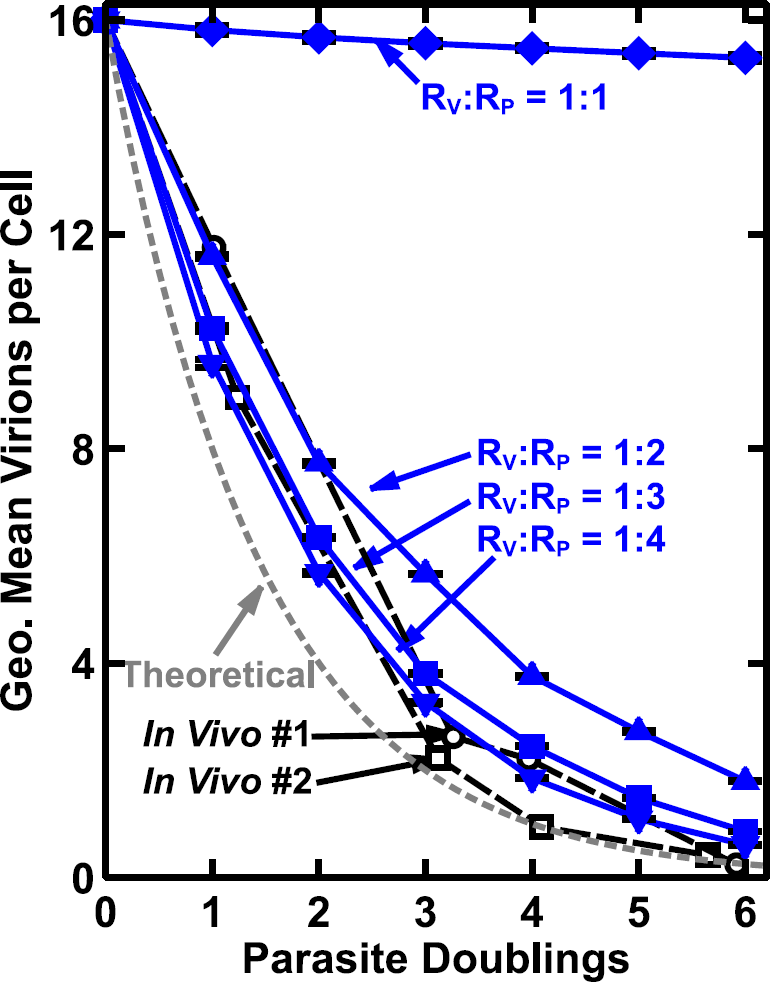
Gillespie simulation of LRV1 loss during 2CMA treatment. Plots of simulated, theoretical and experimental results showing the number of LRV1 virions per cell as a function of cell doubling are shown. Gillespie simulations performed assuming relative inhibition of LRV1 and parasite replication (RV:RP) to be between 1:1 and 1:4 are depicted by blue lines.These lines represent averages and standard deviations of 6 identical simulations (Excel). A theoretical plot is show (gray dotted line) for total inhibition of viral replication with ideal 2-fold viral dilution per population doubling time. Two experimental data sets from Kuhlmann et al. (15) measuring LRV1 loss during 2CMA treatment are shown as black dashed lines.

The effect of 2CMA was then modeled by adjusting the ratio of LRV1 to parasite replication rates (RV:RP). Importantly, at concentrations relevant to these studies, 2CMA has little effect on the growth of the parasite itself (15). After setting RV:RP to 1:2, 1:3, or 1:4, LRV1 was now rapidly lost (Fig. 8). Comparison of the simulations with RV:RP of 1:3 or 1:4 showed close correspondence with the experimentally-measured rate of LRV1 loss under 2CMA inhibition (15) (Fig. 8; solid lines, simulation; dashed lines, experimental).

To compare these predictions with our experimental data on parasite replication and RDRP inhibition, we made an initial simplifying assumption that the primary determinant of LRV1 replication would be the lowest RDRP 2CMA-TP IC50 (150 µM, for full-length product synthesis by LD virions; Table 1; Fig. 3, 4A). Since RDRP inhibition would reduce viral RNA levels within infected cells, RDRP activity would likely become rate-limiting at high 2CMA-TP concentrations. We then calculated the 2CMA-TP concentrations required to give RV:RP of 1:2, 1:3 or 1:4. Importantly, at 10 µM extracellular 2CMA, intracellular 2CMA-TP is 410 µM, and RV:RP was calculated to be ˜1:3 (Fig. 8). Thus the simulation and experimental data are in good agreement about the quantitative and qualitative aspects of RDRP inhibition and viral loss. This suggests that despite the potential complexity of viral replication, the system behaves as though RDRP activity is rate limiting for viral replication and that our *in vitro* measurements of 2CMA-TP inhibition are consistent with our *in vivo* measurements of viral elimination time courses.

## Discussion

Previously, we showed that two 2’-C-methyladenosine analogs selectively inhibit the replication of *Lgy* and *Lbr* LRV1 but have significantly less effect on the parasite growth rates. These compounds were the first such inhibitors to be described for any totivirus (15). There, the mechanism of action was presumed but not shown to follow the classic antiviral nucleoside paradigm of uptake, conversion to the nucleoside triphosphate, and inhibition of the viral RDRP. In this study, we provide evidence that this is in fact the case for *Lgy* LRV1 inhibition following 2CMA treatment. Importantly, the ability of the parasite to accumulate high levels of 2CMA-TP (Fig. 7) was strongly correlated with viral elimination at concentrations above 3 µM external 2CMA (15). These data provide key information for the design of other improved LRV1 inhibitors with greater potential for use as therapeutics.

We first established an assay for RDRP activity from purified virions, separated on CsCl gradients to obtain virions in different stages of maturation (Figs 1-3). RDRP activity was dependent on the presence of LRV1 and yielded two major products, corresponding to the full length viral genome as well as a heterogeneous collection of small and presumably abortive transcripts (Fig. 3). Quantitative analysis showed that the IC50s for full-length product synthesis were somewhat lower than those measured for small product synthesis (p < 0.05), and significantly less for low-density virions than high- (p < 0.001; Table 1). These may reflect the intrinsic sensitivities of the RDRP activity within mature and immature viral particles, perhaps related to the distinct transcriptase/positive-strand (mRNA) and replicase/negative-strand synthesis activities. This is the first such report for Totiviruses, for which antiviral drugs have only recently been reported (15). Differential effects on replicase versus transcriptase activity have also been seen in reoviruses, where ribavirin triphosphate inhibits replicase but not transcriptase activity (42). Our studies were constrained by two factors: first, the virion-based RDRP assay depends on native RNA substrates, and second, the virion-containing fractions used were not homogeneous (Fig. 2). This prevented clean separation of RDRP transcription and replication activities, and precluded the use of tightly-controlled initiation and elongation assays, required for determination of the precise mechanism of RDRP inhibition by 2CMA-TP. Future studies with purified RDRP and defined RNA substrates will be required to fully elucidate the mechanism of action of 2CMA-TP.

2CMA was completely inactive for RDRP inhibition, as were dATP or TTP, structurally-similar non-substrate nucleoside triphosphates (Fig. 5). Activation to triphosphate form is common and often rate limiting amongst nucleoside analog drugs (43-45). It was also shown previously that the triphosphate form of 2CMA, but not the analog itself, is active against the Hepatitis C virus RNA polymerase (46).

Despite the ability of 2CMA to clear LRV1 from parasites grown in micromolar concentrations of drug (15), 2CMA-TP inhibition of LRV1 RDRP activity was not very potent, with IC50s ranging upwards of 150 µM (Table 1). We resolved this discrepancy by showing that the parasites avidly scavenged 2CMA from the medium and efficiently convert it to the active triphosphate form (Fig. 6), reaching intracellular 2CMA-TP concentrations more than 40-fold that of extracellular 2CMA (Fig. 7). This likely reflects the fact that as purine auxotrophs, *Leishmania* parasites must avidly scavenge all naturally occurring purines from their environment, through a combination of powerful transporters and metabolic interconversions (32). One particularly important step for 2CMA and related inhibitors may be adenosine kinase (47,48), which may mediate the initial and often rate-limiting phosphorylation of antiviral nucleosides (44,49). Strong accumulation of toxic anti-leishmanial purines has also been noted in earlier studies (50,51). Thus the salvage pathway converts 2CMA into a potent tool for eliminating the virus.

Importantly, the accumulated levels of intracellular 2CMA-TP closely matched the consequences of RDRP inhibition and LRV1 loss. When grown in 10 µM external 2CMA, internal 2CMA-TP levels greater than 400 µM were attained, well over the minimal LRV1 RDRP IC50 (Table 1). In contrast, at 1 µM external 2CMA, internal 2CMA-TP concentrations were only 80 µM, well below that needed for RDRP inhibition, and little effect was seen on LRV1 levels (15). These experimental observations were corroborated using Gillespie (52) simulation to model LRV1 loss (Fig. 8). Our studies did not examine other potential 2CMA-TP targets such as the capsid endonuclease (53-57) and could not discern the exact mechanism of RDRP inhibition by 2CMA-TP. Nevertheless, these analyses show that the elimination of LRV1 by 2CMA can be largely explained through the direct inhibition of LRV1 RDRP activity by 2CMA-TP.

Previously, we proposed that treatments targeting LRV1 could be used therapeutically to ameliorate the severity of *Lgy* and *Lbr* infections (15,31). 2CMA-TP accumulates rapidly to inhibitory concentrations within 1-2 hr of 2CMA exposure, and once formed, is retained at inhibitory concentrations for >8 hr (Fig. 7). This suggests that a relatively short period of treatment may lead to a prolonged period of viral inhibition. In animal models, 2CMA has a short serum half-life, which precludes its use in humans or animals. The serum half-life of 7d2CMA, however, is markedly longer (0.3 vs. 1.6 hr (58)). Interestingly, 7d2CMA shows an EC50 against Zika virus in cultured mammalian cells of about 10 µM, and dosing regimens have shown significant inhibition in animal models (59,60). This suggests it might likewise be possible to achieve inhibition of LRV1 *in vivo* using 7d2CMA.

In future work, one priority will be the development of anti-LRV1 agents with improved potency. Preliminary studies expressing a promiscuous HSV TK gene within *Leishmania* did not increase the spectrum of activity significantly for those analogs tested from our previous study^2^ (15), suggesting that the lack of activity may reflect failure to inhibit the LRV1 RDRP itself rather than insufficient activation to triphosphates. Similarly, we found that several immucillins shown previously to inhibit *Leishmania* nucleoside hydrolases (Immucillin A, DADMe-Immucillin A, Immucillin H, and DADMe-Immucillin H; (61,62)) had little effect on the potency of 2CMA^2^. Although nucleoside analogs themselves, these immucillins showed no inhibition of LRV1 levels when tested at concentrations up to 100 µM^2^.

Additional priorities will be identifying compounds with more favorable pharmacokinetics and minimal toxicity against the mammalian host cell. Thus, efforts focusing on improved potency against the RDRP activity itself may prove most fruitful. Our work now sets the stage for future studies exploring the possibility that improved LRV1-targeted therapies may ameliorate the pathology of those *Leishmania* species and strains that bear this fascinating virus.

## Experimental Procedures

### Parasite strains and media

Luciferase-expressing isogenic clones of *L. guyanensis* strain M4147 (MHOM/BR/75/M4147) were utilized for these studies. LRV1+ clone LgyM4147/SSU:IR2SAT-LUC(b)c3 and LRV1− clone LgyM4147/pX63HYG/SSU:IR2SATLUC(b)c4 were described previously (63). For some experiments *Lgy* M4147/LRV1+ line expressing GFP+ [LgyM4147/SSU:IR2SATLUC(b)c3/SSU:IR3HYG-GFP+(b)] was used (provided by E. Brettmann). Schneider’s medium (Sigma Aldrich, St. Louis, MO) was prepared following the manufacturer’s instructions, supplemented with 10% heat-inactivated FBS, 0.76 mM hemin, 2 µg/mL biopterin, 50 U/mL penicillin, and 50 µg/mL streptomycin, and adjusted to a final pH of 6.5. M199 medium was prepared with 2% heat-inactivated FBS, 2% filter-sterilized human urine, 0.1 mM adenine, 1 µg/mL biotin, 5 µg/mL hemin, 2 µg/mL biopterin, 50 U/mL penicillin, 50 µg/mL streptomycin, and 40 mM HEPES, pH 7.4 (64). No significant differences were observed in the properties of virus preparations from either medium. Cells were counted using either a hemocytometer or a Coulter counter (Becton Dickinson).

### Purification and fraction of virions on CsCl gradients

Parasites were grown to early stationary phase in M199 or Schneider’s medium (3×10^7^ or 9×10^7^ cells/mL, respectively). 1×10^10^ cells were pelleted at 2200×*g* for 15 min at 4°C and washed twice with 10 mL ice-cold TMN buffer (100 mM Tris, pH 7.5; 50 mM MgCl_2_; 1.5 M NaCl). Cells were then resuspended in 1 mL ice-cold lysis buffer (TMN buffer plus 1 mM DTT, 1× Complete protease inhibitor cocktail (Roche) and 1% (v/v) Triton X-100), homogenized by pipetting 10-12 times with a 1-mL micropipette, and incubated on ice for 20-30 minutes. Lysis was completed by passing the mixture repeatedly through a 27G needle, after which it was clarified by centrifugation at 15,000 × *g* for 10 minutes at 4°C. Density gradients were prepared by thoroughly mixing the clarified lysates with enough 10×TMN buffer, saturated CsCl, and distilled water to make 12 mL of solution at a final density of 1.35 g/mL (2.82 M CsCl). Gradients were spun in a pre-chilled SW41Ti rotor (Beckman) at 32,000 rpm and 4°C for approximately 72 hours. Twelve 1-mL fractions were recovered immediately from each gradient using a density gradient fractionator (ISCO).

The distribution of capsid protein across each gradient was determined by binding of 50 µL aliquots of each fraction to a nitrocellulose membrane using a Mini-fold II Slot-Blot System (Schleicher & Schuell, Keane, NH). The membrane was incubated on a roller with blocking buffer (2% non-fat powdered milk in PBS) for 1 hour, then stained with 1:2500 rabbit anti-capsid antibody (65) in blocking buffer plus 0.2% TWEEN-20 (TWEEN buffer) for another hour. The membrane was then washed 3 times for 5 minutes in 1×PBS plus 0.1% TWEEN-20 (PBST). Membranes were next incubated in TWEEN buffer for 1 hour with 1:10,000 goat anti-rabbit antibodies conjugated to IRDye 680 (LiCor Biosciences). Finally, the membranes were washed 3× in PBST and once in PBS. Membranes were scanned with an Odyssey Infrared Imaging System (LiCor Biosciences). The density of each fraction was measured by taking its refractive index with an Abbe refractometer (Bausch and Lomb) and converting to density using published formulas (66). Gradient fractions of interest (Fig. 2) were dialyzed twice against 1×TMN and once in 1×TMN plus 20% glycerol (4° C), reaching CsCl concentrations less than 2 μM. Fractions were flash frozen and stored at -80°C prior to use.

### RDRP assay

RDRP activity of purified virions was measured using an [α-^32^P]UTP incorporation assay described previously (33). Briefly, 20 µL reactions contained 10 mM Tris-HCl (pH 7.5); 150 mM NaCl; 3 mM MgCl_2_; 4 mM DTT; 50 μM each ATP, CTP, and GTP; 20 µCi [α-^32^P]UTP; and 10 µL virions. Reactions were incubated at room temperature for 1 hour and quenched by addition of 350 µL TRIzol (Ambion). A corresponding gradient fraction from LRV1- parasites was included as a negative control in each set. RNA was purified using a Direct-Zol RNA miniprep kit (Zymo Research) and run on a native 1.2% agarose-TAE gel in a vertical gel apparatus (Owl Scientific) along with dsDNA sizing standards. The standards lane was excised and stained with ethidium bromide, while the radiolabeled products were detected by exposing an imaging plate for 24 hours and reading it with a FLA-5100 phosphoimager (Fuji). The amount of radiolabeled UTP in each RDRP product was quantified using the gel analysis tool in FIJI/ImageJ (67). Equivalent regions from the negative control reaction were also integrated to calculate the background (Fig. 3).

To study inhibition of the viral RDRP by 2CMA-TP, varying amounts of the compound were added to standard RDRP reaction mixtures. 2CMA-TP was custom synthesized by Jena Bioscience, and its identity was confirmed using electrospray ionization with a Fourier-transform mass spectrometer in negative ion mode (Thermo Scientific). The stock concentration of 2CMA-TP was calculated by UV absorption at 260 nm, assuming that its molar extinction coefficient was identical to ATP. To measure the amount of 2CMA-TP which is non-specifically hydrolyzed over the course of an RDRP reaction, mock reactions were run using LRV1- gradient fractions, cold UTP, and 300 μM 2CMA-TP. The 20-µL reactions were diluted to 80 µL with distilled water and immediately analyzed by HPLC as described below.

To estimate IC50 concentrations for each virion population and RDRP product, the quantified RDRP activity data was normalized to untreated and LRV- controls and fitted to logistic dose-response models. Fitting was performed and 95% confidence intervals were estimated using the “drc” package in the R statistical language (68).

### Measurement of parasite volumes

Cultures of WT or GFP-expressing LRV1+ *Lgy* M4147 were seeded at 2×10^5^ cells/mL and analyzed when they reached early, mid, or late log phase. From each sample, one aliquot was analyzed by light scattering on a flow-cytometer, while another was immobilized by treatment with 20 mM sodium azide and imaged by spinning-disk confocal microscopy (69). Cell volumes were calculated using a custom ImageJ script (67) (Supplemental Text 1). A standard curve relating forward scattering intensity to measured volume was developed from this data. Forward scattering intensity of parasites in our assay conditions was then measured and used to estimate the intracellular parasite volume.

### Nucleotide extraction from *Leishmania* parasites

For each sample, 10^8^ cells were collected by centrifugation at 2200 × *g*, 4° C for 5 min, re-suspended in 1 mL ice-cold PBS and re-centrifuged. The cell pellet was gently re-suspended in 100 µL ice-cold 0.5×PBS plus 7 nmol dGTP as a recovery and elution standard. Cells were immediately lysed by rapidly re-suspending in 900 µL ice-cold 5:4 acetonitrile:water mixture (70) and vortexing continuously for 5 min at 4°C. Insoluble debris was pelleted at 16,000 × *g* for 5 minutes and the clarified extract was transferred to a fresh tube. The solvent was removed by evaporation in a Savant SpeedVac concentrator (Thermo Scientific) with the heater off and the vacuum pump refrigeration on. Samples were re-suspended with 80 µL distilled water, flash frozen, and stored at -80°C prior to HPLC analysis.

### HPLC separation of nucleotides

Cell extracts were clarified by centrifuging for 2 min. at 16,000×*g*. Nucleoside di- and tri-phosphates were separated by isocratic HPLC as previously described (71). Briefly, a 20 µL aliquot of clarified extract was injected onto a Zorbax SB-C18 column (5 μm particle size, 250 mm × 4.6 mm, Agilent) and eluted at 1 mL/min with 150 mM KH_2_PO_4_ (pH 6.0); 4.2 mM tetrabutylammonium hydroxide; and 5.4% methanol. Eluting compounds were monitored by UV absorbance at 254 nm (Fig 6). The elution times of nucleoside triphosphates as well as 2CMA and 2CMA-TP were determined by running them individually. A minor peak present in each standard was presumed to represent the di-phosphate form of that nucleoside. A mixture containing 200 μM ATP, GTP, CTP, UTP, and dGTP was used periodically to assess column performance. Peak areas were integrated using Millenium32 software (Waters). Varying amounts of ATP, 2CMA-TP, and dGTP standards were used to construct a calibration curve relating peak areas to compound amounts. Peak areas varied linearly with nucleotide amounts injected down to 50 pmol.

### Gillespie simulation of LRV1 inhibition

We modeled the effects of 2CMA treatment on LRV1 using the next-reaction modification to the Gillespie algorithm (a more detailed description as well as code is provided in Supplemental Text 2) (41). The parameters used to define the system were as follows: number of parasites, number of virions per parasite, parasite growth rate, and virus replication rate. All simulations were initialized with 1000 parasites and 16 virions per cell. Each cell and virus was assigned an amount of time remaining until it divided or replicated, respectively. Because these delay times were composed of an unknown but large number of elementary chemical reactions, they were selected from Gaussian distributions about the mean parasite division and virion replication times, rather than the Poisson distributions used for elementary reactions (41,52). At each step, the event with the shortest time remaining was selected, the simulation time incremented, and the model updated accordingly.

## Acknowledgements

We thank E. Galburt for discussions and advice leading to the Gillespie simulation presented in this work, J. Henderson for assistance developing the HPLC protocol presented here, V. Schramm for providing immucillins, and E. Brettmann for providing GFP-expressing *Lgy*. We also thank A.C.M. Boon, C.E. Cameron, D. Goldberg, P. Olivo, C. Stallings, and N. Tolia for comments and/or suggestions. Supported by NIH grants R01AI029646 and R56AI099364 to SMB and Sigma-Aldrich Predoctoral and Sondra Schlesinger Graduate Student fellowships to JIR.

## Author contributions

JIR and SMB designed the study; JIR performed the experiments; and JIR and SMB analyzed the data, and wrote the paper.

## Footnotes

1 The abbreviations used are: *Leishmania guyanensis* (*Lgy*), *Leishmania braziliensis* (*Lbr*), *Leishmania* RNA virus 1 (LRV1), 2’-C-methyladenosine (2CMA), 2’-Cmethyladenosine triphosphate (2CMA-TP), RNA-dependent RNA polymerase (RDRP), 7-deaza-2’-Cmethyladenosine (7d2CMA), cesium chloride (CsCl), low-density (LD), medium density (MD), high density (HD), phosphate buffered saline (PBS), single stranded RNA (ssRNA), double stranded RNA (dsRNA).

2 J.I.R., unpublished observations.

